# Emotion increases the similarity between neural representations of pictures, but not their perceived similarity

**DOI:** 10.1101/2021.06.21.449164

**Authors:** Martina Riberto, Rony Paz, Gorana Pobric, Deborah Talmi

## Abstract

Stimuli that evoke the same feelings can nevertheless look different and have different semantic meanings. Although we know much about the neural representation of emotion, the neural underpinnings that govern judgements of emotional similarity are unknown. One possibility is that the same brain regions will represent similarity between emotional and neutral stimuli, perhaps with different strengths. Alternatively, emotional similarity could be coded in separate regions, possibly those known to express emotional valence and arousal preferentially. In behaviour, the extent to which people consider similarity along the emotional dimension when they evaluate the overall similarity between stimuli has never been investigated. While the emotional features of stimuli may dominate explicit ratings of similarity, it is also possible that people neglect the emotional dimension as irrelevant. We contrasted these hypotheses with two measures of similarity and two different databases of complex negative and neutral pictures, the second of which afforded exquisite control over semantic and visual attributes. Emotion increased neural similarity in a set of regions that represented both emotional and neutral stimuli, including the inferior temporal cortex, the fusiform face area, and the precuneus. Emotion also increased neural similarity in early visual cortex, anterior insula and dorsal anterior cingulate cortex, despite no increase in BOLD-signal amplitudes in these regions. Despite the stronger neural similarity between emotional stimuli, participants rated pictures taken from two distinct emotional categories as equally similar. These results contribute to our understanding of how emotion is represented within a general conceptual workspace.

## Introduction

We may judge an image of a homeless person and an image of a person injured in a car accident as different, because they have different meanings and probably share very few visual details, or as similar, because both evoke negative feelings. The extent to which similarity along the emotional dimensions influences overall perceived similarity between complex stimuli is not known.

All stimuli can be described according to their location on continuous dimensions, such as valence and intensity, and their proximities reflect aspects of the relationships between them (Russell and Pratt 1980; Chikazoe et al. 2014). This perspective suggests that perhaps stimuli that are entirely neutral, and thus close to the middle of these axes, may be perceived as more similar than stimuli at the extremes. Yet, similarity along specific stimulus dimensions rarely explains more than half the variance in overall similarity perception (Iordan et al. 2018; Dima et al. 2020). Emotional similarity refers to the tendency to group stimuli together because they evoke the same feelings (Riberto, Pobric, and Talmi 2019). It is important to understand emotional similarity because aberrant similarity perception could impact psychological well-being (Puccetti et al. 2021) and be clinically relevant for the overgeneralisation bias in anxiety disorders (Laufer, Israeli, and Paz 2016). Pairwise ratings routinely show higher semantic relatedness among emotional than randomly-selected non-emotional pictures (Talmi 2013), and among positive than negative stimuli (Koch et al. 2016), but in all previous research on similarity perception, stimuli confounded emotional and semantic similarity. Others showed wider generalisation in conditioned than unconditioned stimuli both in healthy controls (Laufer and Paz 2012; Starita et al. 2019) and in anxiety disorders (Ahrens et al. 2016). Research to date, however, has not compared explicit similarity judgements between emotional and neutral stimuli. Our first hypothesis was that emotional similarity will influence explicit ratings of overall similarity, with ratings of emotional stimuli higher than ratings of neutral ones.

Emotional stimuli are thought to be represented in a distributed network of cortical and subcortical regions, which are not only functionally specific to affect (Barrett and Bar 2009), such as the occipitotemporal lobe, which underpins visual-semantic representations of emotional categories (Kragel et al. 2019). Nevertheless, there is agreement that some regions express emotion preferentially, with the insula and the anterior cingulate cortex expressing emotional intensity, and the ventral prefrontal cortex positive valence (Lindquist et al. 2012). Only a handful of studies directly investigated the neural underpinnings of emotional similarity. Multi-voxels pattern analysis (MVPA), such as Representation similarity analysis (RSA), can map how similarity perception is represented in the brain by correlating the data at neural and behavioural level (Kriegeskorte, Mur, and Bandettini 2008). Using this technique, increased neural similarity between exemplars that predicted threat was found in the amygdala (Visser et al. 2013), the occipitotemporal cortex (Dunsmoor et al. 2013) and the superior frontal gyrus (Visser, Scholte, and Kindt 2011). The hippocampus expressed similarity between stimuli associated with reward (Zeithamova et al. 2018) and pain (Wagner, Rütgen, and Lamm 2020), but not emotional pictures (Dandolo and Schwabe 2018). Recent neuroimaging studies found low specificity for each emotion category, and provided evidence against a locationist perspective to the study of emotions (Barrett 2017) reviewed in (Hoemann, Xu, and Barrett 2019). Following from this theory, our second hypothesis was that emotional similarity will be expressed as stronger neural similarity in regions that encode the self-reported participants’ similarity space. Additionally, we expected that regions that express emotional dimensions such as intensity or valence may express emotional similarity uniquely.

We tested these hypotheses in a series of experiments that present several strengths compared to the state-of-the-art. We used different similarity judgements tasks and picture databases, one of which permitted, for the first time, control over taxonomic and thematic similarity, and narrowed our search volume through innovative searchlight approaches.

## Materials and Methods

### Participants

A total of 90 participants were recruited from the University of Manchester (UK), and from the Weizmann Institute of Science (Israel) to take part in the study (age range, 20 –54 years; mean age, 30.14 years; SD, 7.17) (Experiment 1: 20 participants, 10 females; Experiment 2: 40 participants, 20 females; Experiment 3: 29 participants, 12 females; one participant was excluded, because he did not follow the instructions of the task). All participants had normal or corrected-to-normal vision, and were older than 18 years. They gave informed consent prior to the experiment and have been reimbursed for their participation. The exclusion criteria were: a history of neurological (e.g., head injury or concussion) or psychiatric conditions (e.g., depression, anxiety), drug or alcohol abuse, or regular medication that could influence emotional processing. The study was approved by the ethics board of the University of Manchester and of the Weizmann Institute of Science (protocol number 0287–09-TLV).

## Materials

### First database of complex pictures

In experiment 1, we selected 20 images taken from the Nencki Affective Picture System (NAPS) database (Marchewka et al. 2014). Picture IDs that we selected in experiment 1 are reported in Table S1. NAPS has been validated for use in emotional research (Riegel et al. 2016; Wierzba et al. 2015) and consists of 1,356 realistic, high-quality photographs divided into five categories (people, faces, animals, objects, and landscapes). In order to control for visual similarity, we matched the pictures for low-level visual features, that, unlike subjective ratings of visual complexity, are not affected by the arousal complexity bias (Madan et al. 2018) and by the vividness bias (Todd et al. 2013). These measures included the luminance (the average pixel value of the greyscale image) and the contrast (the standard deviation across all the pixels of the greyscale image) (Bex and Makous 2002). In order to quantify the colours within each image, we computed the quantity of red (R), green (G) and blue (B), according to the RGB colour model. Finally, the JPEG size and the entropy of each greyscale image were used as indices of the overall visual complexity of each image (Donderi 2006). The JPEG size was determined with a compression quality setting of 80 (on a scale from 1 to 100). Perceptually simple images are highly compressible and therefore result in smaller file sizes. The entropy, H, is computed from the histogram distribution of the 8-bit grey-level intensity values x: H =–Σp(x)log p(x), where p represents the probability of an intensity value x. H varies with the ‘randomness’ of an image. High-entropy images are noisier and have a high degree of contrast from one pixel to the next, whereas low-entropy images have rather large uniform areas with limited contrast. The sample of images included 10 emotional and 10 neutral images. The designation of images to this category was based on the NAPS ratings of valence and arousal on a 9-points scale provided by 204 European participants. We considered emotional pictures as rated less than 4 in the valence scale (negative valence) and more than 6 in the arousal scale (high arousal), whereas the neutral images ranged from 4 to 6 in both dimensions. To validate the NAPS norms, we also asked our participants to rate the valence and the arousal of the picture before the main task. Table S2 showed the mean and the standard deviation of the different visual and emotional measures for emotional and neutral pictures, as well as the differences between them. We controlled to some extent for semantic similarity by choosing images from the same category - the ‘people’ category. All included more than one person in an outdoor scene. These images contain a lot of information beyond the persons themselves, placing them in a rich and realistic context. The matching we achieved between emotional and neutral conditions exceeds that in most published studies and represents the current state-of-the-art in controlling emotional and neutral stimuli in research. However, the range of emotional themes was reduced compared to that in the neutral set. Therefore, emotional pictures might be rated as more similar, because of the higher thematic similarity compared to neutrals.

### Second database of complex pictures

In experiments 2-3, in order to control the emotional and neutral pictures for thematic similarity, we selected natural scenes in a way that all the categories depicted realistic events that do not co-occur in the environment. In particular, we chose 72 real-world colour photographs using Google images, which represented one or more people in outdoor situations. We divided them into 4 categories according to the scene that was depicted, resulting in 18 images per category. Two of the categories were neutral, and two were emotionally-arousing and negatively valenced. These latter categories represented either poverty scenes (emotional category 1, E1) or car accidents (emotional category 2, E2). The neutral categories portrayed either people hanging laundry to dry (neutral category 1, N1) or talking on the phone (neutral category 2, N2). The full set of pictures can be found at https://dtalmi.wixsite.com/website/resources. We minimised the thematic similarity between emotional categories, by selecting for each of the emotional categories action-context combinations that do not normally occur in a common theme or scenario. The same was true for the two neutral categories. To control for taxonomic similarity to some extent, all the pictures we selected shared two semantic features, they depicted *people outdoor*. Second, we controlled the pictures for affordance, namely the action that a scene can afford, by selecting pictures that depicted only one type of action – and therefore, affordances - in each category. Specifically, in E1, people sit on the ground while begging; in E2, accident victim(s) lay either on a surface (the ground or a crashed car); in N1, people stand hanging and drying clothes; in N2, they stand or walk talking on the phone in the street. Although these actions and affordances differed across the 4 categories, the design ensures that these differences did not influence comparisons across the two neutral and two emotional categories. Finally, we controlled the stimuli for visual properties, as in experiment 1. Finally, an independent sample of 10 healthy participants rated the valence and the arousal of the stimuli. Table S3 showed the mean and standard deviation of visual and emotional measures for each category as well as the differences among them.

### Procedure

A graphical representation of the general experimental procedure is showed in Figure 1. In all the experiments, we asked participants to judge the similarity of a set of complex pictures to test our main hypothesis for the behavioural data: that the perceived similarity between emotional compared to between neutral pictures will be higher. As showed at the top of Figure 1, in the first two experiments participants performed a pairwise similarity rating task. In experiment 1, after rating the valence and arousal of each picture from the first dataset, participants rated all the possible combinations among the stimuli. In experiment 2 we focused on the ratings of interest (denoted with red circles at the bottom of Figure 1), and therefore, participants only rated the similarity between emotional categories (E12) and between neutral categories (N12) of pictures from the second dataset, as well as between emotional and neutral categories (EN), with the latter pairs serving as catch trials. Experiments 1-2 ended after approximately twenty minutes. In experiment 3, after a functional Magnetic Resonance Imaging (fMRI) scan, participants performed a surprise multi-arrangements (MA) task to judge the similarity of the 72 pictures on a bidimensional space, as depicted at the top right of Figure 1 (duration: approximately one hour).

**Figure 1.**
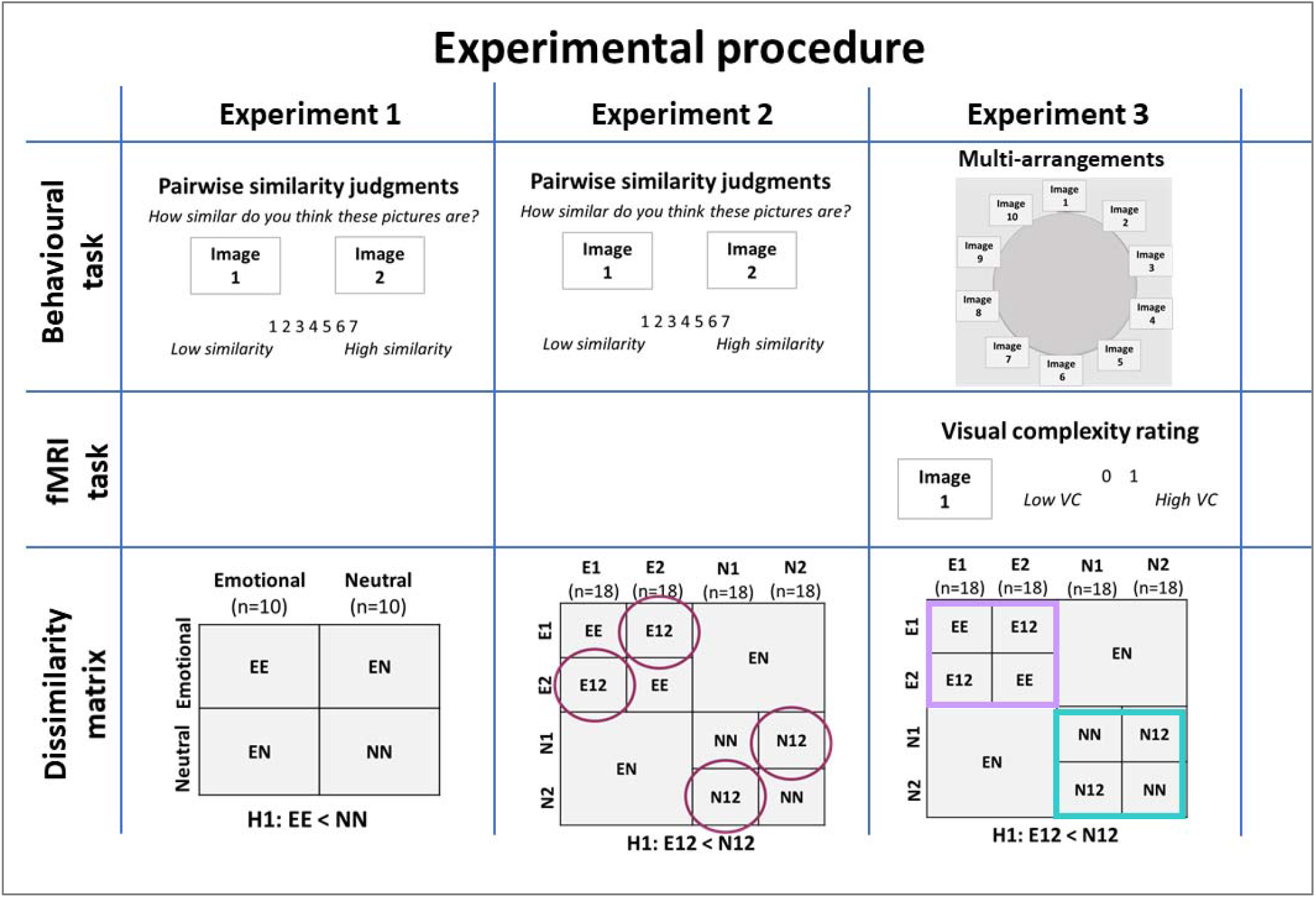
Graphical representation of the experimental procedure. In experiments 1-2, participants performed the same behavioural task. They were presented with a pair of pictures and rated their similarity on a 7-points scale (low to high similarity). In experiment 1, participants judged all the possible combinations from the 1^st^ database, which consisted of 20 complex pictures (10 emotional and 10 neutral) selected from the NAPS. We expected as main finding lower dissimilarity (higher similarity) between emotional (EE) than neutral (NN) pictures. In experiment 2, participants judged the similarity between emotional and neutral pictures from the 2^nd^ database. It consisted of 72 pictures from 4 semantic categories (18 pictures in each category), two emotional (E1 and E2) and two neutral (N1 and N2). Participants only rated E12, N12 and few EN pairs only: E12 represented the similarity between E1 and E2, N12 between N1 and N2, and EN between emotional and neutral pictures. We expected lower dissimilarity (higher similarity) in the former. In both experiments 1-2, EN comparisons served as manipulation checks. The same database was used in experiment 3, wherein participants first judged the subjective visual complexity of each picture during a functional magnetic resonance imaging (fMRI) scan, and then judged the similarity among all the pictures by arranging them in a circular arena. We tested the same hypothesis as in experiment 2, and extended it also to the neural data. The violet square in the dissimilarity matrix represents the ‘emotional similarity space’, and the green one the ‘neutral similarity space’.

### Valence and arousal rating task

To validate the designation of pictures from the two datasets to emotional and neutral conditions, participants completed a valence and arousal rating task, following the procedure suggested by Lang et al. (2008) (Lang, Bradley, and Cuthbert 2008). Each trial started with a central fixation cross for 500 ms. Then, participants viewed one of images presented in the centre of the screen, and rated each pictures on two 9-points scale (valence scale: 1, negative emotions; 9, positive emotions; 5 neutrals. Arousal scale: 1, relaxed; 9, aroused; 5 neutral). We instructed participants to respond as quickly as possible by clicking the appropriate number key, and informed them that there was not a right or wrong answer. Pictures from the first dataset were rated by participants in experiment 1 prior to commencing that experiment, while the ratings of pictures from the second dataset was completed by a separate group of participants.

### Subjective similarity perception

The data from the behavioural experiments were used as subjective measures of similarity, that is, similarity ratings in experiment 1-2, and Euclidean distance in experiment 3.

#### Pairwise similarity rating task

In experiment 1-2, participants rated the similarity of paired pictures on a 7-points scale (1= low similarity, 7= high similarity). In experiment 1, they rated all possible pairwise combinations (190 pairs), resulting from the database of 20 complex pictures. In experiment 2, because of time constraint, we divided the 72 pictures into two subsets (‘even’ and ‘odd’, n=36 within each subsets); in addition, we focused on pairs in E12 and N12, as well as some in EN as catch trials (total pairs= 170; 81 in both E12 and N12, and 8 in EN). We chose the pairwise presentation, because each pair is independently rated and also small differences in similarity judgements can be detected, compared to a triad ‘forced-choice’ similarity task, wherein only binary responses are provided (Goldstone, Medin, and Halberstadt 1997; Miller 1994). We instructed participants to base their judgment on the overall meaning of the picture, without considering any visual details (e.g., the background colour, the number of people), and informed them that there was not a right or wrong answer.

#### Multi-arrangements task

In experiment 3, after the fMRI scan, participants performed a surprise behavioural task, where they were asked to judge the similarity among all the pictures by using the multi-arrangements (MA) task. We chose it because it is a quick and efficient task for acquiring similarity judgements in experiments with a relatively large number of stimuli. Kriegeskorte and Mur (2012) established the MA test-retest reliability (r=0.81) as well as external validity (Kriegeskorte and Mur 2012). The task comprised different trials. In each trial, a subset of 16 stimuli was presented along the perimeter of a circle, or ‘arena’, on a computer screen. Participants had unlimited time to drag and drop the stimuli in the arena according to their similarity, such that similar stimuli were placed close to each other and dissimilar stimuli apart. In other words, the distance among stimuli in the arena reflected their dissimilarity. We instructed participants to focus on the content of the pictures and to ignore visual details (e.g., the colour of the background, the number of people in the scene). A trial ended when participants arranged all the stimuli in the arena. Subsequent trials started with another subset of stimuli to be arranged, selected by using the ‘Lift-the-weakest algorithm for adaptive design of item subsets’. This method optimises trial efficiency by adaptively selecting item subsets whose dissimilarity estimates presented the weakest evidence. The task ended after approximately one hour, when participants judged all the possible combinations among stimuli.

### MRI procedure

In experiment 3, images were acquired on whole body MRI scanner (Trio TIM, Siemens, Germany) with a 12-channel head coil. Functional images were acquired with a susceptibility weighted EPI sequence (TR/TE=2000/30 ms, flip angle=75 degrees, voxel dimensions=3×33×3.5 mm, 192 slices) in 4 separate scanning sessions (up to two minutes between sessions). Anatomical T1-weighted images were acquired after the functional scans (MPRAGE, Repetition time (TR)/Inversion delay time (TI)/Echo time (TE)=2500/900/2.32 ms, flip angle=8 degrees, voxel dimensions=1 mm isotropic, 32 slices).

As showed in Figure 1, during the fMRI scan, participants viewed the 72 complex pictures on a blank screen (size 800 x 800 pixels); we asked them to rate their visual complexity to make them focus on the stimuli, by pressing the right or the left button of the response box, respectively. We instructed participants that there was not a right or wrong answer in the task; rather, they had to focus on their *subjective* perception during the ratings. In order to guide participants in the ratings, we suggested to them that ‘a picture of few objects, colours, or structures would be less complex than a very colourful picture of many objects that is composed of several components’ according to Madan et al. (2018) (Madan et al. 2018). Images were presented in a random order for 3 seconds, during which participants had to make their ratings, interleaved with a black fixation cross (mean jitter 3 seconds). The task was divided into 4 runs, during which every picture was presented once, thus resulting in 4 repetitions for each picture, and a total duration of approximately 50 minutes.

### Data analysis

In the similarity judgements tasks, we expected higher similarity (lower dissimilarity) within-category than between categories. We also expected higher similarity (lower dissimilarity) between emotional than between neutral conditions, as showed at the bottom of Figure 1. The first prediction serves as manipulation check, since a good category boundary simultaneously maximize the within-category similarity, and minimize the between categories similarity; the second prediction represents our main hypothesis, and applies also for the neural data. In experiment 1, EN was calculated by averaging the dissimilarity between emotional and neutral pictures, and the dissimilarity within-emotional (EE) and within-neutral (NN) categories by averaging the dissimilarity between emotional, and between neutral pictures, respectively, for each participant. In experiment 3, EE represented the averaged dissimilarity within E1 and within E2, NN the averaged dissimilarity within N1 and within N2, and EN across both E1 and E2, and N1 and N2, for each participant. Finally, in experiments 2-3, E12 was measured by averaging the dissimilarity between the two emotional categories, and N12 between the two neutral categories. Additional details about the statistical analyses are reported in the following sections.

### Behavioural data analysis

We analysed these data by using Representation similarity analysis (RSA). Specifically, in experiment 1, the similarity ratings were entered as input in a 20 x 20 similarity matrix for each participant. The rows and the columns represented the experimental stimuli, and each cell reflected the similarity rating for each pair. Then, for each subject, a Representational Dissimilarity Matrix (RDM) was computed. We first standardized the similarity ratings, by subtracting 1 (the lowest similarity rating) from each rating *x*, and then divided by 6 (highest similarity rating - lowest similarity rating). Second, we transformed them into correlational distances, by subtracting the ratings from 1. The correlational distance ranges from 0 to 2 (0 for perfect correlation, and thus high similarity; 1 for no correlation; 2 for perfect anticorrelation), and was entered as input in each cell of the RDM. As a consequence, the RDM is symmetric about a diagonal of zeros. Next, we extracted from the single-subject RDM the mean dissimilarity and the standard deviation of the conditions of interest as mentioned in the key hypotheses. These were entered as dependent variables in a repeated-measures ANOVA, with the conditions as grouping factor (experiment 1: EE, NN, and EN; experiment 2: E12, N12, EN). In experiment 3 (MA task), similarity was measured as Euclidean distance between stimuli in the arena. Specifically, at the end of each trial, a *partial* RDM is estimated, showing the Euclidean distance between stimuli within each trial. At the end of the task, a *global* 72 x 72 RDM is estimated by averaging the partial RDMs with an iterative rescaling. This scaling procedure takes into account that in each trial participants focused on a specific subset, and that, therefore, there is not a permanent relationship between screen distance and dissimilarities across trials (see (Kriegeskorte and Mur 2012) for details). Then, we extracted from each participant’s *globa*l RDM the mean and the standard deviation of the conditions of interest mentioned in the section about the key hypotheses. These were entered as dependent variables in 3 repeated-measures ANOVAs. In particular, the formers served to test lower dissimilarity in EE and NN than in EN, and any differences in the dissimilarity in EE and NN. Finally, the latter ANOVA was used to investigate the main hypothesis. Bonferroni post hoc corrections for multiple comparisons (p<0.05) were used to explore the nature of the effect.

We conducted additional analyses to test differences in the variance across participants in the judgements of similarity between emotional than neutral stimuli. With this aim, we conducted two-samples F-tests for variance, one for each contrast of interest: experiment 1: EE vs NN; experiment 2: E12 vs N12; experiment 3: EE vs NN, and E12 vs N12. *Multidimensional scaling* (MDS). We performed the MDS to visualise the structure of the similarity space, wherein proximities reflect similarities among stimuli and are measured on an ordinal scale. The rank order of proximities determines the dimensionality of the space and the metric configuration of the points representing the stimuli (Shinkareva, Wang, and Wedell 2013). As reported in previous studies in this research field, we assumed this space to be bidimensional, with valence and arousal as orthogonal dimensions (Russell and Bullock 1985).The badness-of-fit the MDS representation is estimated with the Stress measure.

#### Analysis of emotional (valence and arousal) and visual complexity ratings

Valence and arousal ratings were entered as dependent variables in two repeated-measures ANOVAs, with picture type (emotional vs neutral) as a within-group factor in experiment 1, and category as a within-group factor in experiment 2-3. Moreover, we analysed the visual complexity ratings from the fMRI task by transforming them into a continuous variable. Specifically, for each subject we calculated the proportion of ‘high complexity’ responses, by dividing the number of ‘high complexity’ responses within each category by 18 (the number of pictures within each category), and then averaged them across sessions. These were entered as dependent variables in a repeated-measure ANOVA, with category (i.e., E1, E2, N1, and N2) as grouping factor. The results from the valence and arousal ratings of experiment 1 are reported in Table S2, those from experiment 2-3 in Table S3 in the supplementary materials. The results of the visual complexity rating task are shown in Table S4. All the results were Bonferroni corrected for multiple comparisons (p<0.05). Data analyses were conducted in Matlab R2018 (MATLAB 2018a, The MathWorks, Inc., Natick, Massachusetts, United States), and SPSS (IBM SPSS Statistics for Windows, Version 25.0. Armonk, NY: IBM Corp).

### Neuroimaging data analysis

#### Preprocessing

Neuroimaging data were pre-processed and analysed using Statistical Parametric Mapping (SPM12) (http://store.elsevier.com/product.jsp?isbn=9780123725608) and MATLAB R2018a (MATLAB 2018a, The MathWorks, Inc., Natick, Massachusetts, United States). Functional images were slice-time corrected to reduce the mismatching between acquisition timing of different slices, and realigned to a reference (mean) image to minimize the variance due to head movements. These were then coregistered to the high-resolution T1-weighted structural image, which was coregistered and normalized to MNI space. Finally, functional images were normalized to a standard template volume based on the Montreal Neurological Institute (MNI) reference brain to achieve a more precise comparison across individuals. Spatial smoothing was performed only on functional data analysed with a conventional univariate approach using a 6-mm full-width at half-maximum isotropic Gaussian kernel. No spatial smoothing was carried over on the multivariate functional data, according to the standard practices for MVPA studies (Kriegeskorte, Mur, and Bandettini 2008; Haxby et al. 2001). After preprocessing, functional data from each voxel were analysed using the general linear model (GLM). Each stimulus was modelled as a separate event beginning with picture presentation onset, using the canonical function in SPM12, and included in the model as regressor of interest (72 regressors per session). Six motion correction parameters were also modelled within each session, and included in the model as regressor of no interest. From this GLM analysis, we obtained a single beta image for each stimulus. Contrast images for each stimulus against the implicit baseline were generated based on the fitted responses, and averaged across sessions. The resulting 72 T-contrast images were used as inputs for RSA. Although our hypotheses were specific to the multivariate representations, we also performed conventional univariate analyses as manipulation check, that is, to demonstrate that the RSA results were not related to differences in the average univariate activations among conditions. The preprocessing for the univariate tests was identical to the one for the RSA with the exception of using a 6-mm FWHM Gaussian smoothing kernel. In addition, for each subject’s GLM, each category was modelled as separate condition (i.e., E1, E2, N1, N2) beginning with each picture presentation onset, using the canonical function in SPM12, and included in the model as regressor of interest (4 regressors per session). From this GLM analysis, we obtained a single beta image for each condition. We then compared emotional and neutral conditions (emotional > neutral), thereby producing one contrasted image for each subject. The contrasted image from each subject was then entered as dependent variable in a one sample t-test. Both the univariate and the multivariate results were inclusively masked to only include our Regions of Interest (ROIs) involved in the visual, semantic and emotional processing of complex pictures, as defined in the paragraph about ROIs definition. In the univariate analysis, no clusters (number of voxels > 10) survived the correction for multiple comparisons.

#### ROIs definition

We defined the ROIs by using the AAL template in WFU Pickatlas toolbox (https://www.nitrc.org/projects/wfu_pickatlas) and Anatomy toolbox (https://www.fil.ion.ucl.ac.uk/spm/ext/#AAL), and constructed with MarsBaR 0.43 (http://marsbar.sourceforge.net). We used WFU_Pickatlas toolbox to define the bilateral early visual cortex (EVC) as Broadmann (Ba) 17, the Dorsomedial Prefrontal cortex (DMPFC) corresponded to the Ba 8 and 9, the Ventromedial Prefrontal cortex (VMPFC) to the Ba 10, and the (dorsal and ventral) Anterior Cingulate Cortex (ACC) to the Ba 32 and 24.The Retrosplenial cortex (RSC), the occipital place area (OPA) and the Parahippocampal place area (PPA) were respectively defined: the bilateral RSC as Ba 29 and Ba 30; the OPA as an 8 mm sphere around the coordinates reported by Julian et al. (2016) (Julian et al. 2016) (left OPA: - 34, -77, 21; right OPA: 34, -77, 21) (Julian et al. 2016); the PPA as an 8 mm sphere around the coordinates reported by (Henson and Mouchlianitis 2007) (left PPA: -27, -45, -12; right PPA 30, -42, -9). The Face Fusiform Area (FFA) was defined as an 8 mm sphere around the coordinates reported by (Henson and Mouchlianitis 2007) (left FFA: -42, -51, -18; right FFA: 42, -45, -21). The medial Temporal lobe (MeTL) comprised the Entorhinal cortex defined with Anatomy toolbox, and the bilateral Hippocampus, the Perirhinal cortex, the Parahippocampal cortex defined with Automated Anatomical Labelling (AAL). The same toolbox was used for the bilateral Inferior Temporal cortex (ITC), the anterior temporal lobe (ATL), the Amygdala, the Thalamus, the Insula, the Precuneus and the bilateral Orbitofrontal cortex (superior, middle, inferior and medial OFC). We combined these ROIs into one ‘ROIs mask’, which was used in the searchlight RSA.

### Quantifying objective (neural) similarity perception

#### Brain- behaviour correlations

In order to test our main hypothesis (i.e., higher neural similarity between the two emotional than the two neutral categories), we first conducted a very precise localisation technique, the searchlight RSA, to investigate which brain regions (within the ROIs mask) represented the participants’ similarity space. This was carried out by computing the Spearman’s correlation between brain activation-patterns RDMs and behavioural RDMs (second order isomorphism). The behavioural RDM represented the participants’ similarity space resulted from the MA task, created as explained in the paragraph about the behavioural data analysis. Three separate analyses were conducted. The first used the entire RDM (with all the 72 stimuli, ‘all RDM’); the second focused exclusively on the emotional stimuli (36 stimuli, ‘emotional RDM’), and the third on the neutral stimuli (36 stimuli, ‘neutral RDM’), depicted as violet and green squares at the bottom of Figure 1, respectively. We conducted these latter two analyses to explore whether any brain region was involved in the representation of either the emotional or the neutral categories. For the purpose of these three analyses, three brain activation-pattern RDMs were constructed for each participant in the same way. The participant’s brain activation-pattern RDMs were computed by entering the T-contrast images into a matrix with all the voxels in the rows, and the experimental stimuli in the columns. Then, for each subject and each of the three analyses, 3 × 3 × 3 voxels spherical cluster was moved throughout the brain and at each location in the ROIs mask a correlational distance (among T values) was assigned to the centre voxel of the sphere, resulting in a (x, y, z, number of pairs) brain activation patterns RDM for each subject. This measure quantified the dissimilarity across voxels in a given searchlight sphere for each specific pair. The number of pairs represented all the possible combinations between experimental stimuli (2556 pairs with 72 stimuli, 630 with 36 stimuli). Next, for each stimulus, the similarity between brain and behavioural RDMs was estimated using a pairwise Spearman’s correlation. This provides a correlational map between the behavioural and the brain RDMs for each subject, which reveals where the similarity space is best represented in the brain (highest correlation), and an ‘n map’, wherein the number of voxels that contributes to each correlational value is reported in each entry. The correlational coefficients were Fisher’s z transformed, and inference was performed at each voxel by performing a one side signed rank test across subjects, testing the null hypothesis of no correlation between brain and behaviour RDMs. The resulting p values (uncorrected) were thresholded to control the false-discovery rate (FDR). We performed two different FDR correction procedures, to yield a more conservative as well as a more lenient set of results. In the conservative procedure, we divided the p values by the total number of voxels in the ROIs mask. In the more lenient procedure, we divided the p values by the number of voxels that contribute to each correlational value (between brain and model RDM). The number of voxels was extracted from the n map associated with the correlational map.

#### Differences in neural dissimilarity between emotional and neutral categories

We conducted a second set of analyses to test our main hypothesis, that is, higher neural similarity (lower dissimilarity) between the two emotional compared to the two neutrals categories (similarity E12>N12). We tested this effect in the brain clusters that we observed to be involved in representing both the entire (72 stimuli) and partial (36 stimuli) participants’ similarity space. With this aim, we created different masks, one for each significant cluster. In case of ROIs that were significantly correlated with both the emotional and the neutral similarity space, we selected the clusters correlated with the neutral similarity space. Then, for each subject and each mask, we computed a brain activation-pattern RDM, where each entry represented the correlational distance (1- Spearman’s correlation) between brain activations across voxels within that mask, and the rows and the columns represented the experimental stimuli. We will refer to this as ROI RDM. It is symmetric about a diagonal of zeros, and resulted in 2556 cells in the lower triangular part that reflected the pairwise dissimilarity of the response patterns associated with the stimuli for each ROI. Then, within each participant ROI RDM, we calculated the mean of the conditions of interest (E12 and N12), and entered them as dependent variables in paired t-tests, one for each cluster (p<0.05). The RSA was performed using the MRC-CBU RSA toolbox for MATLAB (http://www.mrc-cbu.cam.ac.uk/methods-and-resources/toolboxes).

As in the behavioural experiments, we tested any differences in the variance across participants in the neural dissimilarity between E12 and N12. We explored this effect in brain clusters wherein we observed significant differences in neural dissimilarity between E12 and N12. We conducted different two-samples F-tests for variance, one for each cluster.

## Results

In a series of experiments with different datasets of real-world pictures, we explored whether emotions are associated with increased perceived similarity, both subjective (ratings) and objective (neural) similarity. We hypothesised that perceived similarity will be higher (dissimilarity lower) (I) within category, compared to between categories, and (II) between the emotional compared to the neutral stimuli.

### Effect of emotions on subjective similarity perception

Experiment 1 confirmed our hypotheses. We found lower dissimilarity in EE compared to both EN (t (19) =-15.53, p<0.001) and NN (t (19) = -7.10, p<0.001), and in NN than in EN (t (19) =- 5.35, p<0.001). When represented in the similarity space, emotional pictures were displaced closer to each other than neutral pictures. This resulted in a Stress value of 0.05, indicating a good fit of this model. This findings are showed in Figure 2.

**Figure 2.**
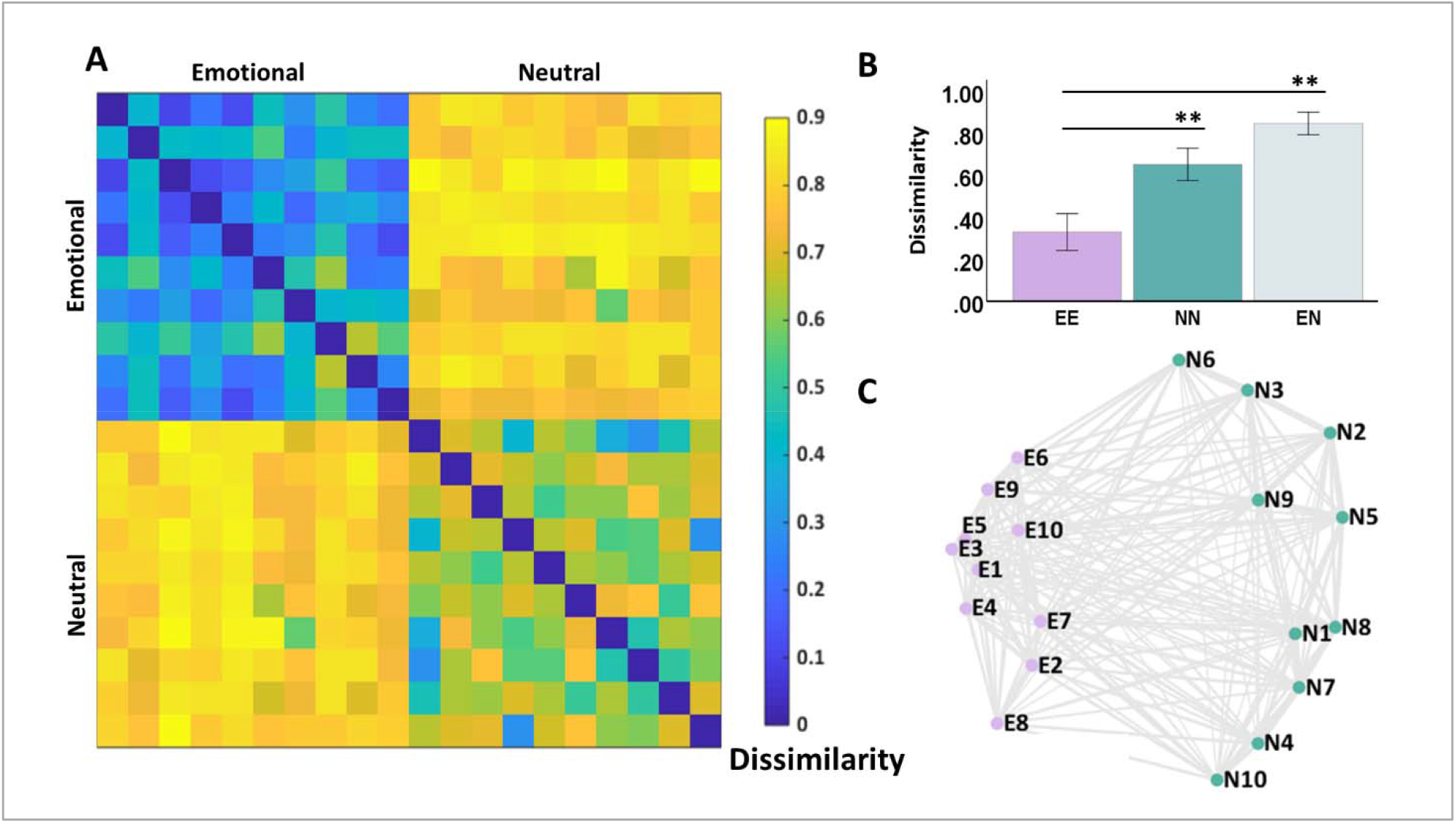
A) Representational Dissimilarity Matrix (RDM) of 20 complex pictures (10 emotional, 10 neutral), averaged across participants. It is symmetric about a diagonal of zeros, the rows and the columns represent the stimuli, and each cell the dissimilarity, measured as 1-standardized similarity ratings between stimuli within each specific pair. Yellow colours denote high dissimilarity, blue colours low dissimilarity. B) The average dissimilarity within emotional pictures (EE), within neutral pictures (NN), and between emotional and neutral pictures (EN, gray). Error bars represent ±2 SEM; **p<0.001. C) The Multidimensional Scaling (MDS) plot of the 20 pictures in a bidimensional space.

In experiment 2, with the second dataset, which controlled for the higher thematic similarity between emotional pictures, we observed different results. Specifically, we found lower dissimilarity in E12 and N12 compared to EN (F (39) = 27.40, p<0.001), but no differences in similarity ratings between the two emotional and the two neutral categories. The same results were replicated using the MA task in experiment 3. We observed lower dissimilarity within category (i.e., EE and NN) than between categories (i.e., E12, N12, EN) (F (29) = 214.76, p<0.001), with no differences in the dissimilarity within categories (F (29) = 4.13, p=0.051), as well as between E12 and N12 (F (29) = 1.55, p=0.22). In the bidimensional space, the proximities between the two emotional, and between the two neutral categories do not differ. The Stress value was 0.10, indicating a fair fit of this model. These findings are showed in Figure 3.

**Figure 3.**
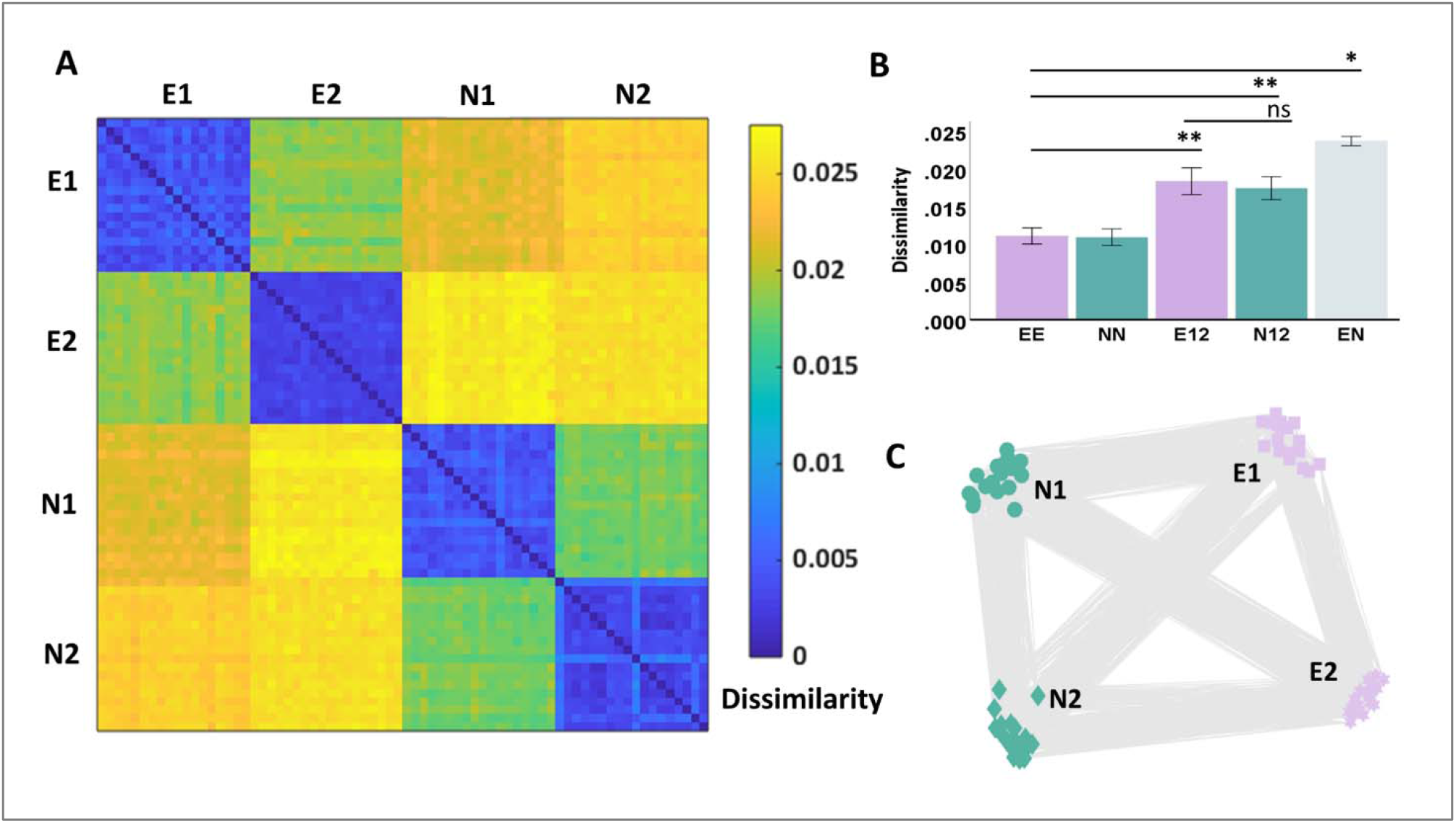
A) Representational Dissimilarity Matrix (RDM) of 72 complex pictures (Emotional categories: E1, poverty (1 to 18); E2, car accidents (19 to 36); Neutral categories: N1, laundry (37 to 54); N2, phone call (55 to 72), averaged across participants. It is symmetric about a diagonal of zeros, the rows and the columns represent the stimuli, and each cell the dissimilarity (measured as Euclidean distance) between stimuli within each specific pair. Yellow colours denote high dissimilarity, blue colours low dissimilarity. B) The average dissimilarity within emotional pictures (averaged across E1 and E2) (EE), within neutral pictures (averaged across N1 and N2) (NN), between emotional pictures (E12), between neutral pictures (N12), and between emotional and neutral pictures (EN). Error bars represent ±2 SEM; *, p_FWE_<0.05; **, p_FWE_<0.001. C) The Multidimensional Scaling (MDS) plot of the 72 pictures in a bidimensional space.

Finally, in all the experiments, we did not observe any significant differences in the variance across participants between emotional and neutral conditions. These results are reported in Table S5.

### Effect of emotions on brain- behaviour correlations

We carried out a searchlight RSA to investigate the brain regions within the ROIs mask that represented the participants’ self-reported similarity space. First, we tested whether the neural-pattern similarity within the ROIs mask was significantly correlated with the entire (72 x 72) similarity space, comprised of neutral and emotional categories. These data only survived our more lenient correction for multiple comparisons (p_FDR_<0.05) (see Materials and Methods). We observed that clusters in the bilateral ITC, the right FFA, and the right Precuneus represented the participants’ similarity space. These findings are reported in in Table 1 and Figure 4A.

**Table 1.** Brain-behaviour correlations. Top: correlations between the entire (72 x 72) stimulus space (named as ‘all RDM’), the and the brain. Significant correlations were observed in the bilateral ITC, right FFA, and the right Prec. Middle: correlations between the emotional (36 x 36) similarity space (named as ‘emotional RDM’) and the brain. Significant correlations were observed in the bilateral EVC, OPA, PPA, FFA, Prec, dACC and left aINS. Bottom: correlations between the neutral (36 x 36) similarity space (named as ‘neutral RDM’) and the brain. Significant correlations were observed in the bilateral OPA, PPA, and left FFA. In all these analyses, correlational coefficients were Fisher’s z transformed, and entered as dependent variables in a one side t test (separately for each brain region), testing the null hypothesis of no correlation between the participants’ similarity space and the neural activation patterns. The resulting p values were thresholded to control for the false-discovery rate (FDR). **, p_FDR_< 0.001. Abbreviations. ITC, Inferior Temporal Cortex; FFA, Face Fusiform Area; Prec, Precuneus; EVC, Early visual cortex; OPA, Occipital place area; PPA, Parahippocampal place area; dACC, dorsal anterior cingulate cortex; aINS, anterior insula. L, Left; R, Right.

| Analysis | Regions | x | y | z | n voxels | t | p<sub>FDR</sub> |
| --- | --- | --- | --- | --- | --- | --- | --- |
| **ALL RDM**<br>**(72 x 72)** | ITC L | 11 | -61 | -9 | 154 | 38.81 | <0.001** |
|  | ITC R | 46 | -63 | -9 | 21 | 40.5 | <0.001** |
|  | FFA R | 42 | -47 | -19 | 69 | 29.64 | <0.001** |
|  | Prec R | 3 | -57 | 20 | 32 | 13.11 | <0.001** |
| **Emotional RDM**<br>**(36 x 36)** | EVC L | -13 | -86 | 4 | 72 | 15.1 | <0.001** |
|  | EVC R | 6 | -93 | -1 | 21 | 16.17 | <0.001** |
|  | OPA L | -34 | -80 | 18 | 103 | 117.25 | <0.001** |
|  | OPA R | 34 | -80 | 18 | 103 | 74.97 | <0.001** |
|  | PPA L | -21 | -43 | -10 | 32 | 58.57 | <0.001** |
|  | PPA R | 24 | -43 | -16 | 29 | 89.19 | <0.001** |
|  | FFA L | -36 | -49 | -22 | 41 | 25.38 | <0.001** |
|  | FFA R | 39 | -49 | -22 | 34 | 26.18 | <0.001** |
|  | Prec | -2 | -60 | 40 | 1249 | 109.58 | <0.001** |
|  | dACC L | -3 | 27 | 1 | 18 | 27.79 | <0.001** |
|  | dACC R | 3 | 15 | 42 | 20 | 29.18 | <0.001** |
|  | aINS L | -35 | 27 | 1 | 18 | 27.79 | <0.001** |
| **Neutral RDM**<br>**(36 x 36)** | OPA L | -29 | -78 | 20 | 11 | 20.44 | <0.001** |
|  | OPA R | 36 | -79 | 16 | 43 | 57.2 | <0.001** |
|  | PPA L | -30 | -49 | -10 | 17 | 18.7 | <0.001** |
|  | PPA R | 25 | -46 | -13 | 10 | 7.19 | <0.001** |
|  | FFA L | -42 | -52 | -13 | 10 | 7.19 | <0.001** |

**Figure 4.**
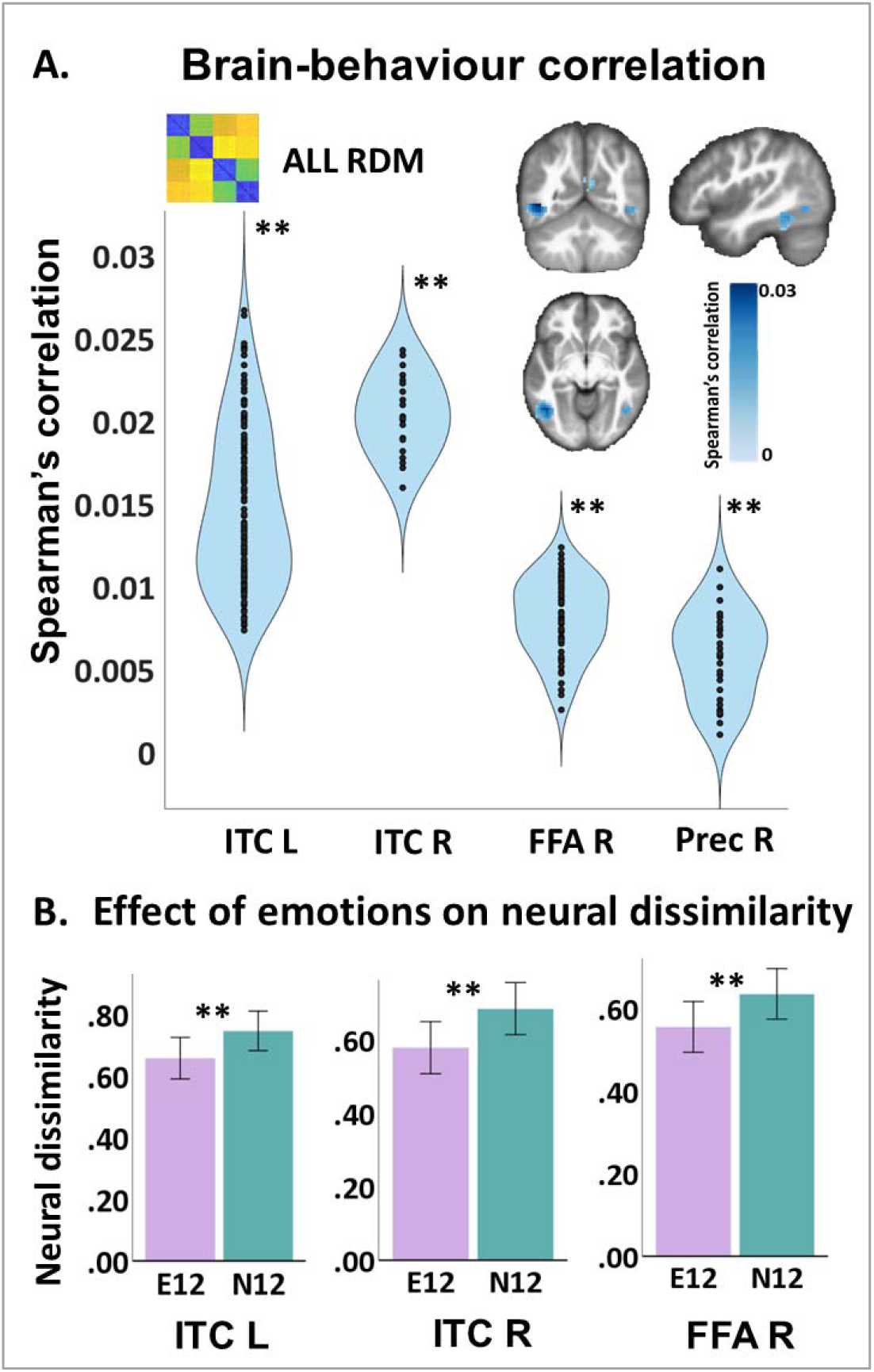
A) Correlation between the entire (72 x 72) stimulus space (named as ‘all RDM’) and the brain. Significant correlations were observed between the behavioural ‘all RDM’ and clusters in the bilateral ITC, right FFA, and the right Prec. Correlational coefficients were Fisher’s z transformed, and entered as dependent variables in a one side t test (separately for each brain region), testing the null hypothesis of no correlation between the participants’ similarity space and the neural activation patterns. The resulting p values were thresholded to control for the false-discovery rate (FDR). **, p_FDR_< 0.001. B) Differences in neural dissimilarity (measured as correlational distance) between emotional and neutral stimuli in different brain clusters, including the bilateral ITL, and the right FFA. The dissimilarity between emotional categories (E12) was calculated by averaging the dissimilarity between E1 and E2, and the dissimilarity between neutral categories (N12) by averaging the dissimilarity between N1 and N2, for each participant. These were entered as dependent variables in paired t tests, one for each brain cluster (p<0.05). **, p< 0.001. Abbreviations. ITC, Inferior Temporal Cortex; FFA, Face Fusiform Area; L, Left; R, Right.

Second, we performed the same analysis separately for the emotional and neutral pictures to explore whether any brain region was involved in the representation of either the emotional or the neutral categories (see the violet and green squares in Figure 1). The results from these analyses survived the more conservative correction for multiple comparisons (p_FDR_<0.05) (see Materials and Methods). We found that participants’ emotional similarity space was significantly correlated with clusters in lower and higher-level visual processing regions, as well as regions involved in emotional processing. These included the bilateral EVC, bilateral OPA, bilateral PPA, bilateral FFA, bilateral Precuneus, bilateral dorsal ACC and in the left anterior Insula. By contrast, participants’ neutral similarity space was significantly correlated with clusters in higher-level visual regions only, including the bilateral OPA, the bilateral PPA, and left FFA. These findings are reported in Table 1 and Figure 5A and 6A.

**Figure 5.**
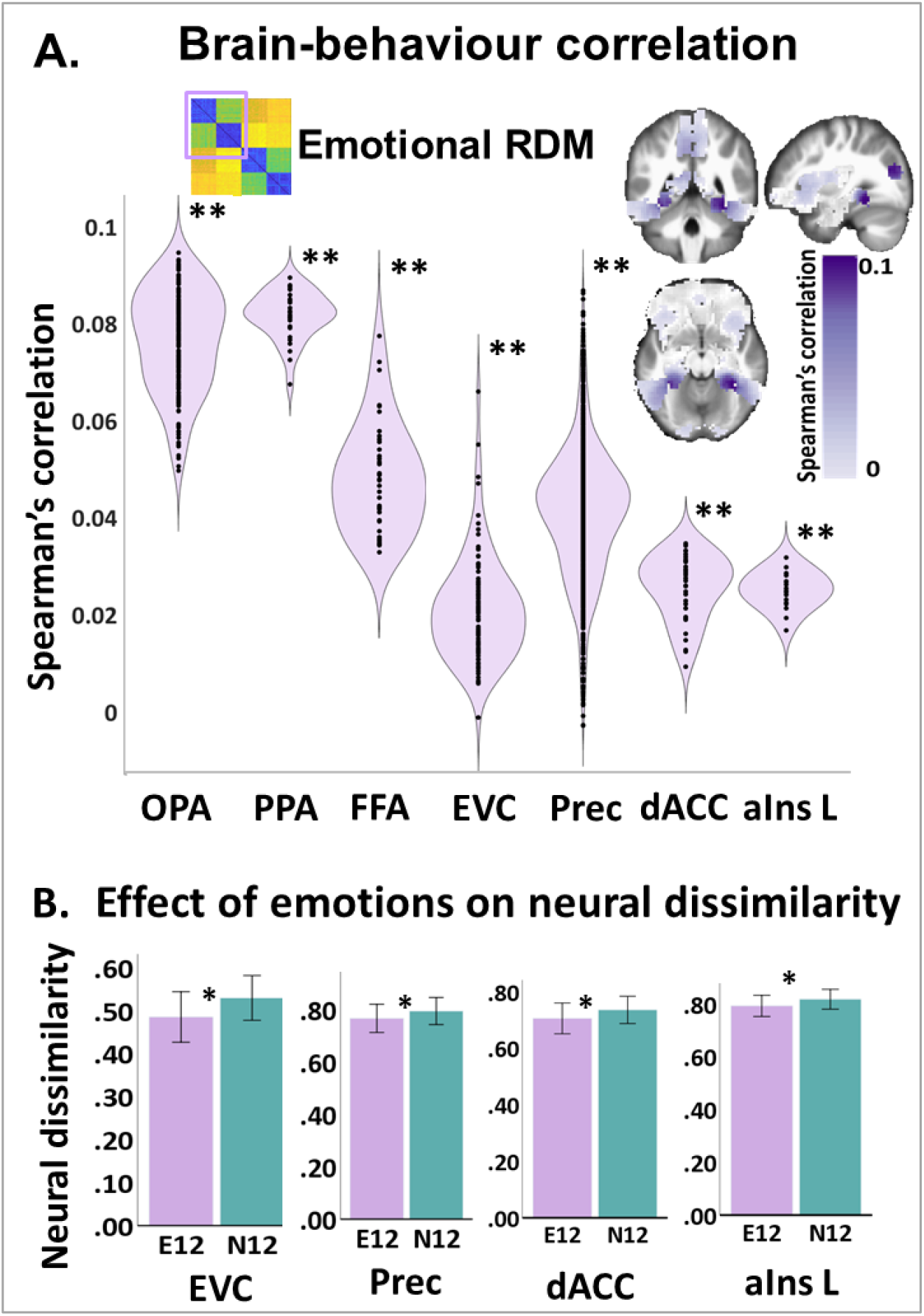
A) Correlation between the emotional (36 x 36) similarity space (named as ‘emotional RDM’) and the brain. Significant correlations were observed between the behavioural ‘emotional RDM’ and clusters in the bilateral OPA, PPA, FFA, EVC, Prec, dACC, and left aIns. Correlational coefficients were Fisher’s z transformed, and entered as dependent variables in a one side t test (separately for each brain region). For simplicity, we averaged the left and the right sides of the clusters wherein both sides were significant. The resulting p values were thresholded to control the false-discovery rate (FDR). **, p_FDR_< 0.001. B) Differences in neural dissimilarity (measured as correlational distance) between emotional and neutral stimuli in different brain clusters, including the bilateral EVC, Prec, dACC and left aIns. The dissimilarity between emotional categories (E12) was calculated by averaging the dissimilarity between E1 and E2, and the dissimilarity between neutral categories (N12) by averaging the dissimilarity between N1 and N2, for each participant. These were entered as dependent variables in paired t tests, one for each brain cluster (p<0.05). *, p< 0.05. Abbreviations. OPA, Occipital place area; PPA, Parahippocampal place area; FFA, Face fusiform area; EVC, Early visual cortex; Prec, Precuneus; dACC, Dorsal anterior cingulate cortex; aIns, Anterior insula; L, left; E12, dissimilarity between emotional categories; N12, dissimilarity between neutral categories.

**Figure 6.**
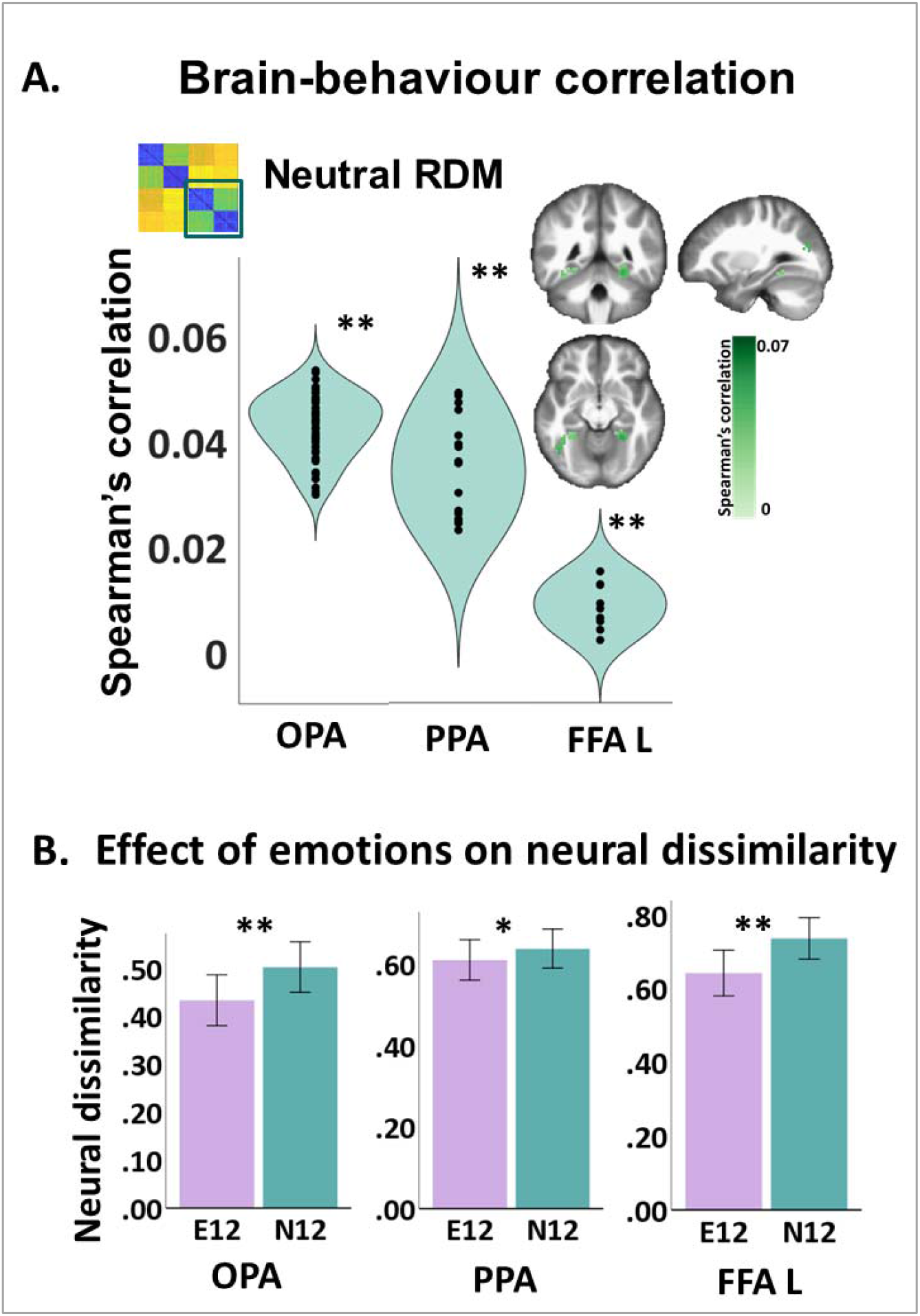
A) Correlation between the neutral (36 x 36) similarity space (named as ‘neutral RDM’) and the brain. Significant correlations were observed between the behavioural ‘neutral RDM’ and clusters in in the bilateral OPA, PPA and left FFA. Correlational coefficients were Fisher’s z transformed, and entered as dependent variables in a one side t test (separately for each brain region). For simplicity, we averaged the left and the right sides of the clusters when both sides were significant. The resulting p values were thresholded to control the false-discovery rate (FDR). *, p_FDR_< 0.05; **, p_FDR_< 0.001. B) Differences in neural dissimilarity (measured as correlational distance) between emotional and neutral stimuli in different brain clusters, including the bilateral OPA, PPA and left FFA. The dissimilarity between emotional categories (E12) was calculated by averaging the dissimilarity between E1 and E2, and the dissimilarity between neutral categories (N12) by averaging the dissimilarity between N1 and N2, for each participant. These were entered as dependent variables in paired t tests, one for each brain cluster (p<0.05). *, p< 0.05; **, p< 0.001. Abbreviations. OPA, Occipital place area; PPA, Parahippocampal place area; FFA, Face fusiform area; L, left; E12, dissimilarity between emotional categories; N12, dissimilarity between neutral categories.

### Effect of emotions on objective (neural) similarity

We performed the ROIs RSA to explore whether the neural representations of emotional categories are more similar than those associated with neutral categories. This analysis was carried out in brain clusters from the above analysis, namely, those that significantly correlated with the whole participants’ similarity space (Figure 4B), as well as with its emotional (Figure 5B), and neutral (Figure 6B) similarity spaces. As predicted, the neural pattern dissimilarity of emotional categories was lower than the one of neutral stimuli in all the previously reported clusters (p<0.05), apart for the right PPA. In addition, we observed trends towards significance in support to our hypothesis in the right EVC (p=0.11) and in one cluster in the left PPA (p=0.06). These findings are reported in Table 2 and in Figures 4B, 5B and 6B.

**Table 2.**
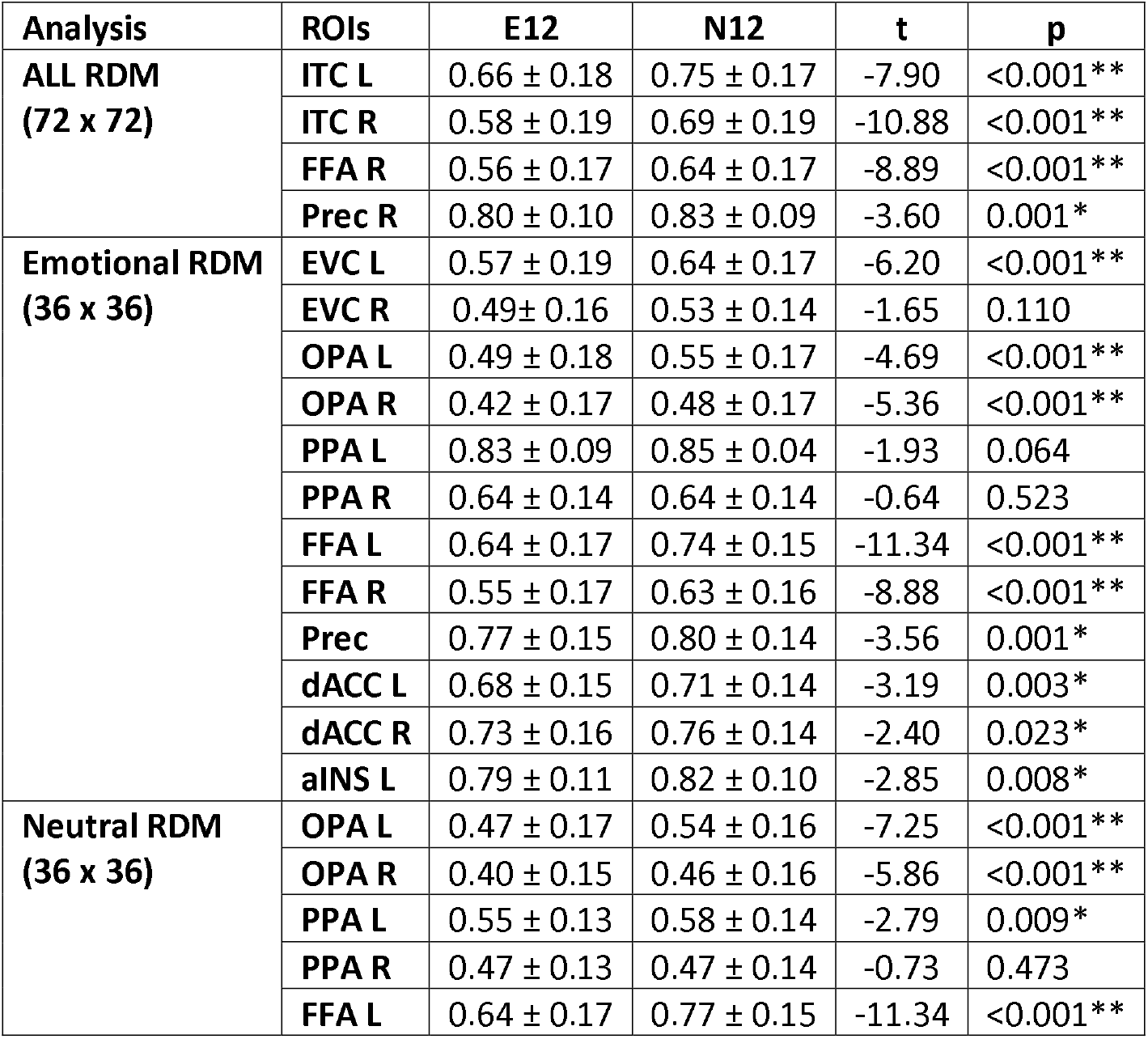
Effect of emotions on neural dissimilarity. Difference in neural dissimilarity (measured as correlational distance) among conditions. The dissimilarity between emotional categories (E12) was calculated by averaging the dissimilarity between E1 and E2, and the dissimilarity between neutral categories (N12) by averaging the dissimilarity between N1 and N2, for each participant. These measures were first computed in brain clusters significantly involved in the representation of the whole (72 stimuli) participants’ similarity space (top of the table). Then, we computed E12 and N12 in brain clusters significantly involved in the representation of the emotional (middle of the table) and neutral (bottom of the table) participants’ similarity space. We entered them as dependent variables in paired t tests, one for each brain cluster. Bonferroni post hoc corrections for multiple comparisons (p<0.05) are summarized at the bottom. *, p_FWE_< 0.05; **, p_FWE_< 0.001. Abbreviations. E12, neural dissimilarity between emotional categories; N12, neural dissimilarity between neutral categories. ITC, Inferior temporal cortex; FFA, Face fusiform area; Prec, Precuneus; EVC, Early visual cortex; OPA, Occipital place area; PPA, Parahippocampal place area; dACC, Dorsal anterior cingulate cortex; AI, Anterior insula; L, left; R, Right.

Finally, we did not observe any significant differences in the variance across participants between E12 and N12 in any brain clusters. These results are reported in Table S6.

## Discussion

We investigated the effect of negative emotions on the perception of similarity between complex stimuli and on the objective similarity between their neural representations using two different similarity judgements tasks and databases of stimuli, the second of which was extremely well controlled. We report two novel findings. First, participants subjectively perceived stimuli from two negatively-valenced, emotionally-arousing categories to be just as similar as stimuli from the two neutral categories, evident in their similarity judgements. Second, the similarity among the neural representations of stimuli from the two emotional categories was higher compared to the neutral ones despite no increase in BOLD-signal amplitudes. This effect was obtained while participants were processing individual stimuli, without any demand to process inter-stimulus relationships, and despite extremely high level of control over visual and semantic stimulus attributes. Some, but not all, of the clusters which expressed similarity among emotional stimuli preferentially also expressed similarity among neutral stimuli. Taken together, these results demonstrated a decoupling between subjective and objective measures of similarity. We discuss the implications of these results below.

### Negative emotion does not increase the subjective similarity between pictures of realistic events

In experiment 1, when thematic similarity was not controlled, we found higher similarity between emotional, compared to neutral, stimuli. In a bidimensional valence-arousal space, emotional stimuli were placed closer to each other than neutral ones. The goodness of fit of the model was good, suggesting that affective features were the most salient in similarity judgements. This result is in keeping with dimensional perspectives on emotions (Barrett and Russell 1999) and recent empirical data (Cowen and Keltner 2017), although our results cannot distinguish between effects based on valence or arousal dimensions. Strikingly, when we controlled for the typically higher thematic relatedness between emotional stimuli in experiments 2-3, by selecting stimuli from four separate semantic categories (two emotional and two neutral), stimuli from the two emotional categories were perceived as equally similar to those from the neutral categories. They were clustered according to the four categories, and the goodness of fit of the bidimensional space dropped to fair, pointing to the semantic meaning of each picture – not negative emotion - as the most relevant feature in ratings of similarity.

The pattern of findings across the three experiments suggests that participants’ conceptual workspace comprises of several integrated dimensions (Prince and Konkle 2020), which include not only perceptual and affective, but also semantic, features. These features may obtain different weights, or relevance, in the subjective perception of similarity according to the relationships among the experimental stimuli (Todd et al. 2020), in agreement with previous literature attesting to the strong context effects on similarity ratings (Goldstone, Medin, and Halberstadt 1997). By including four separate categories in experiments 2-3, we may have increased the weight of the dimension of semantic meaning in overall subjective similarity perception compared to experiment 1, where stimuli were not grouped by category. By contrast, in experiment 1, by using a random selection of both emotionally negative-arousing and neutral pictures, the weight of the affective feature may have increased so that it became the most salient. The relative contribution of semantic and affective features to the overall subjective perception of similarity could be tested in future by manipulating the weight of the semantic and emotional dimensions through task instructions or a mood manipulation. This might have implications for semantic cognition research, which so far have not integrated the relative contribution of affective and semantic dimensions to semantic categorisation Lambon Ralph 2014).

An alternative explanation of our results is that evaluating the similarity between emotional pictures can be affected by individuals’ skill in emotional differentiation, also named emotional granularity (Barrett et al. 2001). High-granular individuals would be more aware of the differences of their emotional experiences when viewing pictures from the two emotional categories used here and may therefore rate them as less similar, while low-granular individuals may rate them as more similar, ultimately masking the difference between emotional and neutral categories. However, if this explanation was correct, we would expect increased variance in ratings of emotional pictures. Instead, there were no significant differences in rating variance between emotional and neutral categories.

### Negative emotion increased the objective similarity between neural representations of realistic events

In experiment 3, we observed stronger neural similarity among stimuli from the two emotional than the two neutral categories, in brain clusters involved in encoding participants’ entire similarity space. They encompassed the ventral visual stream, potential neurobiological underpinning of semantic processing and categorisation (Iordan et al. 2015; Clarke and Tyler 2014), and brain regions involved in affect representations (i.e., precuneus and anterior insula) (Kim, Shinkareva, and Wedell 2017) and modulation (i.e., dorsal anterior cingulate cortex)(Saarimäki et al. 2018). To our knowledge, this is the first report of the neural underpinnings of emotional similarity for a pictures set controlled for visual and semantic attributes, extending findings of Levine et al. (2018) about neural representations of idiosyncratic affective space in the insula (Levine et al. 2018). Our ability to reveal this dissociation may have stemmed from the constrained search volume we used to test our hypothesis regarding neural similarity (rather than through a whole brain search), and from the control exerted here on visual properties of the pictures.

These finding may also have implications for research about the neurobiological correlates of semantic categorisation and generalisation. Previous studies (Dunsmoor et al. 2013; Visser, Scholte, and Kindt 2011) observed an increase in neural similarity among exemplars that predicted threat. They proposed that this mechanism was adaptive, because it enables individuals to differentiate emotionally salient stimuli from those that are not, and might support broad generalisation between physically dissimilar items, which predict the same fitness-relevant outcome. Although our work differed from these studies, where the emotional response was induced though Pavlovian conditioning, we found the same effect here. This converging evidence might suggest that it is evolutionarily more important to integrate the emotional information in neural representations in order to increase the relevance and generalisability of stimuli that might predict a negative outcome. These findings concur with the conclusions of Todd et al. (2020), that emotion serves as a fundamental feature of cognition: it acts as a filter by defining what is bad or good for us, such that any representation of the world is an integrated product between emotion, perception and thought (e.g., “That is a good thing”) rather than discrete and isolated psychological events (e.g., “That is a thing. I feel good”). Our behavioural results suggest that the evolved sensitivity to emotion may be dampened when the context suggests it is less relevant.

In this study, we extended previous findings about the brain regions involved in representing emotional categories and dimensions by exploring, for the first time, differences in the neural representations of the relationships among and between emotional and neutral stimuli. In particular, we observed greater neural similarity between the two emotional than the two neutral categories in the bilateral ITC, right FFA and right precuneus, which were involved in representing the entire similarity space. As part of the hierarchical network located in the ventral visual stream, the ITC integrates relevant low and high level features, resulting into an emergent category structure (Prince and Konkle 2020). Accumulating research agreed on inferior occipitotemporal regions as potential neurobiological underpinnings of semantic categorisation and perceived similarity of objects (Kriegeskorte et al. 2008; Iordan et al. 2015), faces (Guntupalli, Wheeler, and Gobbini 2016; Haxby et al. 2011) and places (Groen et al. 2018; Epstein and Baker 2019). In particular, as showed by Groen et al. (2018), other regions in the ITC involved in action observation and in representing ‘acting bodies’, including FFA, take part to scenes encoding. Accordingly, Brooks et al. (2019) demonstrated that the single subjects’ conceptual space predicts the neural pattern activation in the right FFA (Brooks et al. 2019). We observed stronger neural similarity between emotional categories in these regions (Kim, Shinkareva, and Wedell 2017), probably because of the influence of the precuneus, which is involved in valence representation and structurally connected with the ITC (Lin et al. 2020).

When we investigated the emotional and the neutral parts of participants’ similarity space, we observed higher emotional similarity also in the EVC, OPA and PPA, as well as in the dACC and anterior insula. As aforementioned, OPA and PPA relate low-level visual features encoded in the EVC with the high-level aspects of the scene (Epstein and Baker 2019), and like other ITC regions, may be modulated by regions that are sensitive to emotion, such as anterior insula and dACC (Lindquist et al. 2012), resulting in higher similarity between emotional than neutral regions. Interestingly, our finding that the insula represented emotional, but not neutral, similarity replicate those of Levine and colleagues (2018), who reported that neural similarity there represented emotional similarity, although they did not control their stimulus set for semantic similarity. Finally, the same effect was observed in the EVC, which relies on more fine-grained representations of the stimuli (high granularity) (Coutanche, Solomon, and Thompson-Schill 2016), and encodes low-level visual features of the stimuli that afford the decoding of a broad range of emotions categories (Petro et al. 2013; Barrett and Bar 2009). Specific combinations of low-level visual features (e.g., local colour, luminance, contrast) along with high-level information (e.g., presence of faces, scenes or objects) can act as cues and afford specific categories of emotional response that are more salient, and thus defined, than neutral categories (Kragel et al. 2019).

We also expected that other ROIs, including the orbitofrontal, ventral and dorsomedial prefrontal cortex, were involved in representing the entire similarity space or just the emotional part. However, we did not find any significant correlations with the behavioural data there, perhaps due to the implicit processing of affect in experiment 3. Nor did we observe significant correlations with the amygdala, perhaps because of the strong habituation of amygdala response to repeated stimuli (Plichta et al. 2014).

## Limitations

Our study presents several limitations that can be addressed in future works. First, our materials were limited. We studied only negative emotions, and were able to include only two categories within each level of affect. We chose this to increase the statistical power in terms of number of trials per condition, while keeping the experiment short enough to ensure participants attention. It would be interesting in future studies to examine whether the functional dissociation we observed is replicated with a richer set of materials, including a greater number of semantic categories, and examine whether the greater neural similarity reflects similarity along the dimensions of both valence and arousal, or sensitive to discrete emotional categories (e.g. fear or disgust-inducing stimuli). Second, the experimental stimuli were presented in a random order during a rapid event related design. This is a common procedure in neuroimaging literature. However, it might have had an effect on our results by increasing the trial-by-trial similarity (increased correlation), since overlapping BOLD signals cannot be decorrelated unless multiple trials are combined into a single regressor (Visser et al. 2016). This could have the effect of decreasing our statistical power. Finally, we cannot infer any causal role of emotions on neural similarity from our study. Future studies could use TMS to further explore this aspect of the findings.

## Conclusion

In conclusion, stimuli that share negative feelings are perceived as more similar to each other unless care is taken to remove shared taxonomic and thematic links between them. Once thematic links are controlled, negative emotion does not always increase perceived similarity among realistic events. A set of brain regions expressed emotional similarity preferentially. They included regions that encoded the semantic meaning of the stimuli, as well as those which could modulate them. These results suggest that emotional stimuli are encoded in a more similar fashion not only in brain clusters that are functionally specific to affect, probably because it is evolutionary advantageous for the brain to represent affect more broadly.

Taken together, we observed an intriguing dissociation, for the first time, between subjective similarity, expressed in explicit ratings, and objective similarity, expressed in correlations between neural representations of single stimuli, where only the latter were sensitive to negative emotion. This could be due to dynamically changing weights of the multiple dimensions of participants’ conceptual workspace.

## Supporting information

SI_new

## References

Ahrens, Lea M, Paul Pauli, Andreas Reif, Andreas Mühlberger, Gernot Langs, Tim Aalderink, and Matthias J Wieser. 2016. ‘Fear conditioning and stimulus generalization in patients with social anxiety disorder’, Journal of Anxiety Disorders, 44: 36–46.

Barrett, Lisa Feldman. 2017. ‘The theory of constructed emotion: an active inference account of interoception and categorization’, Social cognitive and affective neuroscience, 12: 1–23.

Barrett, Lisa Feldman, and Moshe Bar. 2009. ‘See it with feeling: affective predictions during object perception’, Philosophical Transactions of the Royal Society B: Biological Sciences, 364: 1325–34.

Barrett, Lisa Feldman, James Gross, Tamlin Conner Christensen, and Michael Benvenuto. 2001. ’Knowing what you’re feeling and knowing what to do about it: Mapping the relation between emotion differentiation and emotion regulation’, Cognition & Emotion, 15: 713–24.

Barrett, Lisa Feldman, and James A Russell. 1999. ‘The structure of current affect: Controversies and emerging consensus’, Current directions in psychological science, 8: 10–14.

Bex, Peter J, and Walter Makous. 2002. ‘Spatial frequency, phase, and the contrast of natural images’, JOSA A, 19: 1096–106.

Brooks, Jeffrey A, Junichi Chikazoe, Norihiro Sadato, and Jonathan B Freeman. 2019. ‘The neural representation of facial-emotion categories reflects conceptual structure’, Proceedings of the National Academy of Sciences, 116: 15861–70.

Chikazoe, Junichi, Daniel H Lee, Nikolaus Kriegeskorte, and Adam K Anderson. 2014. ‘Population coding of affect across stimuli, modalities and individuals’, Nature neuroscience, 17: 1114.

Clarke, Alex, and Lorraine K Tyler. 2014. ‘Object-specific semantic coding in human perirhinal cortex’, Journal of Neuroscience, 34: 4766–75.

Coutanche, Marc N, Sarah H Solomon, and Sharon L Thompson-Schill. 2016. ‘A meta-analysis of fMRI decoding: Quantifying influences on human visual population codes’, Neuropsychologia, 82: 134–41.

Cowen, Alan S, and Dacher Keltner. 2017. ‘Self-report captures 27 distinct categories of emotion bridged by continuous gradients’, Proceedings of the National Academy of Sciences, 114: E7900–E09.

Dandolo, Lisa C, and Lars Schwabe. 2018. ‘Time-dependent memory transformation along the hippocampal anterior–posterior axis’, Nature communications, 9: 1–11.

Dima, Diana C, Tyler Tomita, Christopher Honey, and Leyla Isik. 2020. ‘The representational space of action perception’, Journal of Vision, 20: 1161–61.

Donderi, Don C. 2006. ‘Visual complexity: a review’, Psychological bulletin, 132: 73.

Dunsmoor, Joseph E, Philip A Kragel, Alex Martin, and Kevin S LaBar. 2013. ‘Aversive learning modulates cortical representations of object categories’, Cerebral Cortex, 24: 2859–72.

Epstein, Russell A, and Chris I Baker. 2019. ‘Scene perception in the human brain’.

Goldstone, Robert L, Douglas L Medin, and Jamin Halberstadt. 1997. ‘Similarity in context’, Memory & Cognition, 25: 237–55.

Groen, Iris IA, Michelle R Greene, Christopher Baldassano, Li Fei-Fei, Diane M Beck, and Chris I Baker. 2018. ‘Distinct contributions of functional and deep neural network features to representational similarity of scenes in human brain and behavior’, Elife, 7: e32962.

Guntupalli, J Swaroop, Kelsey G Wheeler, and M Ida Gobbini. 2016. ‘Disentangling the representation of identity from head view along the human face processing pathway’, Cerebral Cortex, 27: 46–53.

Haxby, James V, M Ida Gobbini, Maura L Furey, Alumit Ishai, Jennifer L Schouten, and Pietro Pietrini. 2001. ‘Distributed and overlapping representations of faces and objects in ventral temporal cortex’, Science, 293: 2425–30.

Haxby, James V, J Swaroop Guntupalli, Andrew C Connolly, Yaroslav O Halchenko, Bryan R Conroy, M Ida Gobbini, Michael Hanke, and Peter J Ramadge. 2011. ‘A common, high-dimensional model of the representational space in human ventral temporal cortex’, Neuron, 72: 404–16.

Henson, Richard N, and Elias Mouchlianitis. 2007. ‘Effect of spatial attention on stimulus-specific haemodynamic repetition effects’, NeuroImage, 35: 1317–29.

Hoemann, Katie, Fei Xu, and Lisa Feldman Barrett. 2019. ‘Emotion words, emotion concepts, and emotional development in children: A constructionist hypothesis’, Developmental psychology, 55: 1830.

Iordan, Marius Catalin, Cameron T Ellis, Michael Lesnick, Daniel N Osherson, and Jonathan Cohen. 2018. “Feature Ratings and Empirical Dimension-Specific Similarity Explain Distinct Aspects of Semantic Similarity Judgments.” In CogSci.

Iordan, Marius Cătălin, Michelle R Greene, Diane M Beck, and Li Fei-Fei. 2015. ‘Basic level category structure emerges gradually across human ventral visual cortex’, Journal of cognitive neuroscience, 27: 1427–46.

Julian, Joshua B, Jack Ryan, Roy H Hamilton, and Russell A Epstein. 2016. ‘The occipital place area is causally involved in representing environmental boundaries during navigation’, Current Biology, 26: 1104–09.

Kim, Jongwan, Svetlana V Shinkareva, and Douglas H Wedell. 2017. ‘Representations of modality-general valence for videos and music derived from fMRI data’, NeuroImage, 148: 42–54.

Koch, Alex, Hans Alves, Tobias Krüger, and Christian Unkelbach. 2016. ’A general valence asymmetry in similarity: Good is more alike than bad’, Journal of Experimental Psychology: Learning, Memory, and Cognition, 42: 1171.

Kragel, Philip A, Marianne C Reddan, Kevin S LaBar, and Tor D Wager. 2019. ‘Emotion schemas are embedded in the human visual system’, Science advances, 5: eaaw4358.

Kriegeskorte, Nikolaus, and Marieke Mur. 2012. ‘Inverse MDS: Inferring dissimilarity structure from multiple item arrangements’, Frontiers in psychology, 3: 245.

Kriegeskorte, Nikolaus, Marieke Mur, and Peter A Bandettini. 2008. ‘Representational similarity analysis-connecting the branches of systems neuroscience’, Frontiers in systems neuroscience, 2: 4.

Kriegeskorte, Nikolaus, Marieke Mur, Douglas A Ruff, Roozbeh Kiani, Jerzy Bodurka, Hossein Esteky, Keiji Tanaka, and Peter A Bandettini. 2008. ‘Matching categorical object representations in inferior temporal cortex of man and monkey’, Neuron, 60: 1126–41.

Lambon Ralph, Matthew A. 2014. ‘Neurocognitive insights on conceptual knowledge and its breakdown’, Philosophical Transactions of the Royal Society B: Biological Sciences, 369: 20120392.

Lang, Peter J, Margaret M Bradley, and Bruce N Cuthbert. 2008. “International affective picture system (IAPS): affective ratings of pictures and instruction manual. University of Florida, Gainesville.” In.: Tech Rep A-8.

Laufer, Offir, David Israeli, and Rony Paz. 2016. ‘Behavioral and neural mechanisms of overgeneralization in anxiety’, Current Biology, 26: 713–22.

Laufer, Offir, and Rony Paz. 2012. ‘Monetary loss alters perceptual thresholds and compromises future decisions via amygdala and prefrontal networks’, Journal of Neuroscience, 32: 6304–11.

Levine, Seth M, Anja Wackerle, Rainer Rupprecht, and Jens V Schwarzbach. 2018. ‘The neural representation of an individualized relational affective space’, Neuropsychologia, 120: 35–42.

Lin, Yueh-Hsin, Isabella M Young, Andrew K Conner, Chad A Glenn, Arpan R Chakraborty, Cameron E Nix, Michael Y Bai, Vukshitha Dhanaraj, R Dineth Fonseka, and Jorge Hormovas. 2020. ‘Anatomy and White Matter Connections of the Inferior Temporal Gyrus’, World Neurosurgery, 143: e656–e66.

Lindquist, Kristen A, Tor D Wager, Hedy Kober, Eliza Bliss-Moreau, and Lisa Feldman Barrett. 2012. ‘The brain basis of emotion: a meta-analytic review’, Behavioral and brain sciences, 35: 121–43.

Madan, Christopher R, Janine Bayer, Matthias Gamer, Tina B Lonsdorf, and Tobias Sommer. 2018. ‘Visual complexity and affect: ratings reflect more than meets the eye’, Frontiers in psychology, 8: 2368.

Marchewka, Artur, Łukasz Żurawski, Katarzyna Jednoróg, and Anna Grabowska. 2014. ‘The Nencki Affective Picture System (NAPS): Introduction to a novel, standardized, wide-range, high-quality, realistic picture database’, Behavior research methods, 46: 596–610.

Miller, George A. 1994. ‘The magical number seven, plus or minus two: Some limits on our capacity for processing information’, Psychological review, 101: 343.

Petro, Lucy S, Fraser W Smith, Philippe G Schyns, and Lars Muckli. 2013. ‘Decoding face categories in diagnostic subregions of primary visual cortex’, European Journal of Neuroscience, 37: 1130–39.

Plichta, Michael M, Oliver Grimm, Katrin Morgen, Daniela Mier, Carina Sauer, Leila Haddad, Heike Tost, Christine Esslinger, Peter Kirsch, and Adam J Schwarz. 2014. ‘Amygdala habituation: a reliable fMRI phenotype’, NeuroImage, 103: 383–90.

Prince, Jacob S, and Talia Konkle. 2020. ‘Computational evidence for integrated rather than specialized feature tuning in category-selective regions’, Journal of Vision, 20: 1577–77.

Puccetti, Nikki A, Stacey M Schaefer, Carien M Van Reekum, Anthony D Ong, David M Almeida, Carol D Ryff, Richard J Davidson, and Aaron S Heller. 2021. ‘Linking amygdala persistence to real-world emotional experience and psychological well-being’, Journal of Neuroscience, 41: 3721–30.

Riberto, Martina, Gorana Pobric, and Deborah Talmi. 2019. ‘The emotional facet of subjective and neural indices of similarity’, Brain topography, 32: 956–64.

Riegel, Monika, Łukasz Żurawski, Małgorzata Wierzba, Abnoss Moslehi, Łukasz Klocek, Marko Horvat, Anna Grabowska, Jarosław Michałowski, Katarzyna Jednoróg, and Artur Marchewka. 2016. ‘Characterization of the Nencki Affective Picture System by discrete emotional categories (NAPS BE)’, Behavior research methods, 48: 600–12.

Russell, James A, and Merry Bullock. 1985. ‘Multidimensional scaling of emotional facial expressions: similarity from preschoolers to adults’, Journal of Personality and Social Psychology, 48: 1290.

Russell, James A, and Geraldine Pratt. 1980. ‘A description of the affective quality attributed to environments’, Journal of personality and social psychology, 38: 311.

Saarimäki, Heini, Lara Farzaneh Ejtehadian, Enrico Glerean, Iiro P Jääskeläinen, Patrik Vuilleumier, Mikko Sams, and Lauri Nummenmaa. 2018. ‘Distributed affective space represents multiple emotion categories across the human brain’, Social cognitive and affective neuroscience, 13: 471–82.

Shinkareva, Svetlana V, Jing Wang, and Douglas H Wedell. 2013. ‘Examining similarity structure: multidimensional scaling and related approaches in neuroimaging’, Computational and mathematical methods in medicine, 2013.

Starita, Francesca, Marijn CW Kroes, Lila Davachi, Elizabeth A Phelps, and Joseph E Dunsmoor. 2019. ‘Threat learning promotes generalization of episodic memory’, Journal of Experimental Psychology: General, 148: 1426.

Talmi, Deborah. 2013. ‘Enhanced emotional memory: Cognitive and neural mechanisms’, Current Directions in Psychological Science, 22: 430–36.

Todd, Rebecca M, Vladimir Miskovic, Junichi Chikazoe, and Adam K Anderson. 2020. ‘Emotional objectivity: Neural representations of emotions and their Interaction with cognition’, Annual review of psychology, 71: 25–48.

Todd, Rebecca M, Taylor W Schmitz, Josh Susskind, and Adam K Anderson. 2013. ‘Shared neural substrates of emotionally enhanced perceptual and mnemonic vividness’, Frontiers in behavioral neuroscience, 7: 40.

Visser, Renée M, Michelle IC de Haan, Tinka Beemsterboer, Pia Haver, Merel Kindt, and H Steven Scholte. 2016. ‘Quantifying learning-dependent changes in the brain: Single-trial multivoxel pattern analysis requires slow event-related fMRI’, Psychophysiology, 53: 1117–27.

Visser, Renée M, H Steven Scholte, Tinka Beemsterboer, and Merel Kindt. 2013. ‘Neural pattern similarity predicts long-term fear memory’, Nature neuroscience, 16: 388–90.

Visser, Renée M, H Steven Scholte, and Merel Kindt. 2011. ‘Associative learning increases trial-by-trial similarity of BOLD-MRI patterns’, Journal of Neuroscience, 31: 12021–28.

Wagner, Isabella C, Markus Rütgen, and Claus Lamm. 2020. ‘Pattern similarity and connectivity of hippocampal-neocortical regions support empathy for pain’, Social cognitive and affective neuroscience, 15: 273–84.

Wierzba, Małgorzata, Monika Riegel, Anna Pucz, Zuzanna Leśniewska, Wojciech Łukasz Dragan, Mateusz Gola, Katarzyna Jednoróg, and Artur Marchewka. 2015. ‘Erotic subset for the Nencki Affective Picture System (NAPS ERO): cross-sexual comparison study’, Frontiers in psychology, 6: 1336.

Zeithamova, Dagmar, Bernard D Gelman, Lea Frank, and Alison R Preston. 2018. ‘Abstract representation of prospective reward in the hippocampus’, Journal of Neuroscience, 38: 10093–101.

