## Supplementary material for "Emotion increases the similarity between neural representations of pictures, but not their perceived similarity": SI_new

| **Emotional** | | **Neutral** | |
| --- | --- | --- | --- |
| 205 | 016 | 104 | 165 |
| 127 | 038 | 100 | 035 |
| 226 | 075 | 150 | 095 |
| 238 | 007 | 066 | 249 |
| 235 | 022 | 146 | 057 |

|  |  | **Categories** | | **Statistics** | |
| --- | --- | --- | --- | --- | --- |
|  |  | **Emotional** | **Neutral** | **t** | **p** |
| **Visual measures** | **Luminance** | 89.42 ± 27.31 | 11.13 ± 36.06 | -1.45 | .16 |
|  | **Contrast** | 61.97 ± 1.16 | 62.19 ± 1.14 | -.49 | .96 |
|  | **R** | 93.24 ± 26.06 | 111.62 ± 26.67 | -1.48 | .17 |
|  | **G** | 88.99 ± 27.92 | 110.12 ± 37.84 | -1.41 | .19 |
|  | **B** | 82.58 ± 26.54 | 105.95 ± 41.24 | -1.56 | .15 |
|  | **JPEG** | 337121.90 ± 90579.45 | 27707.50 ± 69785.15 | 1.66 | .11 |
|  | **Entropy** | 7.50 ± .28 | 7.49 ± .24 | .11 | .91 |
| **Emotional measures** | **Valence** | 1.985 ± .743 | 4.840 ± .976 | -10.22 | .000** |
|  | **Arousal** | 7.170 ± 1.389 | 4.840 ± 1.219 | 8.93 | .000** |

**Table S1.** Picture IDs from the NAPS database (‘people’ category), divided into emotional and neutrals (experiment 1).

**Table S2.** Differences in visual and emotional measures between emotional (n=10) and neutral (n=10) pictures (experiment 1). The mean and the standard deviation of each measure are shown, as well as the t and the p value associated with each difference. **, p_FWE_< 0.001.

|  |  | Categories | | | | Statistics | |
| --- | --- | --- | --- | --- | --- | --- | --- |
|  |  | **E1** | **E2** | **N1** | **N2** | **F** | **p** |
| Visual  measures | **Luminance** | 127.43 ± 27.26 | 116.17 ± 29.44 | 124.81 ± 29.28 | 122.11 ± 12.95 | .66 | .580 |
|  | **Contrast** | 63.30 ± 11.40 | 67.16 ± 7.63 | 69.34 ± 1.75 | 69.57 ± 1.33 | 1.36 | .269 |
|  | **R** | 137.89 ± 27.91 | 119.60 ± 29.18 | 133.70 ± 32.52 | 125.01 ± 12.24 | 1.79 | .171 |
|  | **G** | 124.05 ± 27.52 | 114.49 ± 29.90 | 122.90 ± 29.26 | 121.32 ± 14.00 | .51 | .654 |
|  | **B** | 117.38 ± 27.66 | 115.82 ± 3.58 | 111.35 ± 29.07 | 118.51 ± 18.37 | .25 | .829 |
|  | **Jpeg** | 88395.39 ± 1286.54 | 90291.06 ± 1508.73 | 83717.67± 1399.74 | 7864.11 ± 23072.68 | 1.61 | .212 |
|  | **Entropy** | 7.65 ± .22 | 7.68 ± .15 | 7.62 ± .25 | 7.66 ± .16 | .26 | .841 |
| Emotional measures | **Valence** | 2.91 ± 1.42 | 1.97 ± 1.02 | 4.91 ± .26 | 5.13 ± .30 | 46.93 | 000** |
|  | **Arousal** | 6.64 ± 1.40 | 7.74 ± 1.34 | 4.72 ± 1.37 | 4.53 ± 1.26 | 27.37 | 000** |

**Table S3.** Differences in visual and emotional measures among categories. The mean and the standard deviation of each measure are shown, as well as the F and the p value associated with each difference. Abbreviations: E1, Emotional category 1 (poverty scenes, n=18); E2, Emotional category 2 (car accidents, n=18); N1, neutral category 1 (laundry scenes, n=18); N2, neutral category 2 (telephone scenes, n=18) (experiment 2-3).

| Categories | | | | Statistics | |
| --- | --- | --- | --- | --- | --- |
| E1 | **E2** | **N1** | **N2** | **F** | **p** |
| .308 ± .163 | .728 ± .211 | .297 ± .187 | .267 ± .170 | 5.34 | <.001** |
| Post hoc | | | | | |
| E1 vs E2 | **E1 vs N1** | **E1 vs N2** | **E2 vs N1** | **E2 vs N2** | **N1vs N2** |
| -.42, .000** | .01,.1.00 | .04,.1.00 | .43,.000** | .46,.000** | .03,1.00 |

**Table S4.** Differences in visual complexity ratings among categories. The proportion of high complexity ratings within each category (total number of ‘high complexity’ responses divided by 18) was averaged across sessions. Mean and standard deviation of each category, and the statistics of the difference among them are reported at the top of the table. Bonferroni post hoc corrections for multiple comparisons (p<0.05) are summarized at the bottom. *, p_FWE_< 0.05; **, p_FWE_< 0.001.

| **Behavioural experiments** | **Conditions** | | **F value** | **F critical** | **P value** |
| --- | --- | --- | --- | --- | --- |
| **Experiment 1** | EE  0.04 | NN  0.03 | 1.28 | 2.53 | 0.36 |
| **Experiment 2** | E12 | N12 | 0.72 | 1.89 | 0.16 |
| **Experiment 3** | EE  0.000 | NN  0.000 | 0.79 | 2.13 | 1.45 |
|  | E12  0.000 | N12  0.000 | 1.34 | 2.13 | 0.44 |

**Table S5.** Differences in the variance in similarity judgements between emotional and neutral stimuli. The variance averaged across participants for each conditions, and the statistics of each difference between conditions are reported. In experiment 1, EE and NN represent the variance within emotional, and within neutral stimuli, respectively, averaged across participants. In experiment 2-3, E12 and N12 signify the variance between the two emotional, and the two neutral categories, respectively, averaged across participants. Finally, in experiment 3, EE and NN represent the variance within E1 and E2, and within N1 and N2, averaged across participants.

| **ROIs** | **E12** | **N12** | **F value** | **F critical** | **P value** |
| --- | --- | --- | --- | --- | --- |
| **ITC** | 0.03` | 0.03 | 1.04 | 2.13 | 0.45 |
| **Precuneus** | 0.02 | 0.02 | 1.09 | 2.13 | 0.82 |
| **EVC** | 0.02 | 0.02 | 1.27 | 2.13 | 0.52 |
| **OPA** | 0.02 | 0.02 | 1.02 | 2.13 | 0.97 |
| **PPA** | 0.02 | 0.02 | 0.99 | 2.13 | 1.00 |
| **FFA** | 0.02 | 0.02 | 1.07 | 2.13 | 0.85 |
| **dACC** | 0.02 | 0.02 | 1.28 | 2.13 | 0.52 |
| **aIns L** | 0.01 | 0.01 | 1.15 | 2.13 | 0.71 |

**Table S6.** Differences across participants in the variance in neural dissimilarity between emotional and neutral stimuli. The variance averaged across participants for each conditions within each cluster, and the statistics of each difference between conditions are shown. E12 and N12 represent the variance between the two emotional, and the two neutral categories, respectively, averaged across participants. For simplicity, we averaged the left and the right sides of the clusters. Abbreviations. ITC, Inferior Temporal Cortex; EVC, Early visual cortex; OPA, Occipital place area; PPA, Parahippocampal place area; FFA, Face fusiform area; Prec, Precuneus; dACC, Dorsal anterior cingulate cortex; aIns, Anterior insula; L, left; E12, dissimilarity between emotional categories; N12, dissimilarity between neutral categories.
